# Nascent polypeptide within the exit tunnel ensures continuous translation elongation by stabilizing the translating ribosome

**DOI:** 10.1101/2021.02.02.429294

**Authors:** Yuhei Chadani, Nobuyuki Sugata, Tatsuya Niwa, Yosuke Ito, Shintaro Iwasaki, Hideki Taguchi

## Abstract

Continuous translation elongation, irrespective of amino acid sequences, is a prerequisite for living organisms to produce their proteomes. However, the risk of elongation abortion is concealed within nascent polypeptide products. Negatively charged sequences with occasional intermittent prolines, termed intrinsic ribosome destabilization (IRD) sequences, destabilizes the translating ribosomal complex. Thus, some nascent chain sequences lead to premature translation cessation. Here, we show that the risk of IRD is maximal at the N-terminal regions of proteins encoded by dozens of *Escherichia coli* genes. In contrast, most potential IRD sequences in the middle of open reading frames remain cryptic. We found two elements in nascent chains that counteract IRD: the nascent polypeptide itself that spans the exit tunnel and its bulky amino acid residues that occupy the tunnel entrance region. Thus, nascent polypeptide products have a built-in ability to ensure elongation continuity by serving as a bridge and thus by protecting the large and small ribosomal subunits from dissociation.

## Introduction

The ribosome is the central player in translation, at the heart of the central dogma, in all organisms. Accurate and continuous peptide bond formation by the ribosome ensures the uniformity and quality of cellular proteins. Otherwise, aberrant polypeptides would lead to undesired outcomes such as impaired or dysregulated protein functions and aggregation, which perturb cellular homeostasis. Thus, living organisms have multilayered surveillance systems over ribosomes and translation, including proofreading of codon-anticodon recognition (Ieong et al., 2016; Noel and Whitford, 2016; Rodnina and Wintermeyer, 2001), quality control pathways to sense unusual translation reactions, and degradation of premature products (Buskirk and Green, 2017; Inada, 2017; Keiler, 2008).

Translation is initiated when the small ribosomal subunit (the 30S in prokaryotes) recognizes the start codon within mRNA, and this is followed by the recruitment of the initiator aminoacyl-tRNA and the association of the large ribosomal subunit (50S in prokaryotes). In the decoding process, aminoacyl-tRNAs are delivered as ternary complexes with EF-Tu into the A-site of the ribosome complex. After the decoding, EF-Tu dissociates from the ribosome. The peptidyl transferase center (PTC) of the ribosome catalyzes peptide bond formation between the P-site and the A-site amino acids. The ribosomal subunits then dynamically rotate to translocate the peptidyl-tRNA through the A/P hybrid state to the P/P classic state, with the concomitant one-codon advancement of the ribosome along the mRNA. The complex is now ready to proceed to the next cycle of peptidyl-transfer for continuous translation elongation. The nascent peptide grows until a release factor recognizes the stop codons within the A-site and then cleaves the peptidyl-tRNA ester bond. Once the synthesized polypeptide is released, the ribosome is split into two subunits by the cooperative actions of the ribosome recycling factor and EF-G to recycle them for the next round of translation (Choi et al., 2015; Rodnina, 2016, 2018; Rodnina and Wintermeyer, 2016).

Structural studies have invariably shown that the ribosomes possess a tunnel structure (exit tunnel) that accommodates 30-40 amino acid-long segments of tRNA-linked nascent polypeptides (Jenni and Ban, 2003; Voss et al., 2006). During the elongation process, each amino acid residue of a growing nascent polypeptide travels from the PTC through the exit tunnel of the large ribosomal subunit to the extra-ribosomal space (Wilson and Beckmann, 2011). The tunnel is composed of the large ribosomal RNA and the tips of ribosomal proteins, such as uL4, uL22, and uL23 (Nissen et al., 2000). In contrast to the initial suggestion that the tunnel interior is inert in terms of interactions with the nascent polypeptide (Nissen et al., 2000), in reality, the tunnel interacts with nascent polypeptide chains, thus contributing to biological regulation (Chiba et al., 2009; Gong and Yanofsky, 2002; Ishii et al., 2015; Nakatogawa and Ito, 2002; Onouchi et al., 2005; Yanagitani et al., 2011, Ito and Chiba, 2013). Cryo-EM structures of ribosomes stalled by ribosome arresting peptides (RAPs) have revealed a range of interactions between the nascent chains and the ribosomal interior components at the PTC and/or the exit tunnel, such as the uL22/uL4 constriction site (Bhushan et al., 2011; Seidelt et al., 2009; Shanmuganathan et al., 2019; Sohmen et al., 2015; Su et al., 2017). Recent analyses using ribosome profiling (Dao Duc and Song, 2018; Han et al., 2020; Requião et al., 2016; Sabi and Tuller, 2015) and direct detection of accumulating peptidyl-tRNAs (Chadani et al., 2016) have suggested the widespread occurrence of translation pausing, to which nascent polypeptides may contribute. Thus, nascent polypeptides are not simple intermediates in translation; in contrast, they actively regulate the ribosomal functions.

Our recent study demonstrated that nascent polypeptides within the ribosome can induce intrinsic ribosome destabilization (IRD), leading to a premature cessation of translation (Chadani et al., 2017). When the ribosome translates a sequence with consecutive negatively-charged residues or intermittent with prolines (DE/P motif), its subunit structure becomes destabilized. Such nascent polypeptides become susceptible to peptidyl-tRNA hydrolase (Pth), which otherwise cannot access a substrate sequestered within the intact ribosome complex (Das and Varshney, 2006; Sharma et al., 2014). Thus, translation is aborted stochastically within the DE/P motif. Deleting bL31, which bridges the large and small subunits, destabilizes the 70S structure and enhances IRD. Thus, specific amino acid sequences within a nascent polypeptide can compromise the ribosome’s structural integrity and translation continuity. Remarkably, IRD contributes to gene regulation. MgtL, the leader peptide of MgtA (Gall et al., 2016; Park et al., 2010; Zhao et al., 2011), an Mg^2+^ transporter, contains a potential IRD-inducing motif, EPDP, at its N-terminus. The IRD in MgtL is exaggerated under ribosome-destabilizing conditions, including lowered Mg^2+^ concentrations, triggering the MgtA-upregulating genetic scheme. Whereas MgtL provides an example of programmed IRD in cells, our knowledge about the roles of nascent peptides in regulating the stability, in either direction, of the ribosomes in action remains incomplete.

Here, we report our proteome-wide profiling of the potential IRD-inducing sequences (hereafter, called “IRD sequences”) in *E. coli* and our analyses of how this organism maintains robust translation continuity by overcoming IRD. We identified dozens of N-terminal IRD sequences that cause significant premature translation cessation. By contrast, the IRD-like sequences in the open reading frames’ internal regions are silent, raising the possibility that a preceding (N-terminal) nascent polypeptide segment residing in the exit tunnel antagonizes IRD. We found two IRD-counteracting elements in nascent polypeptides: their lengths that occupy the tunnel and the bulkiness of the amino acid side chains in the PTC-adjacent region. The “length” and “bulkiness” of nascent chains affect the genome-wide amino acid distribution in ORFs to minimize the risk of nonproductive translation discontinuation. These results shed new light on the pivotal roles played by nascent chains in the ribosomal exit tunnel in ensuring the efficient synthesis, without delay, of every possible peptide with any sequence context required by living organisms.

## Results

### N-terminal DE/P-rich sequences in *E. coli* ORFs induce IRD

We previously showed that the “EPDP” sequence following the initiator methionine of *E. coli* MgtL destabilizes the translating 70S ribosome from within to prematurely abort translation, which is harnessed as an environmental sensing strategy (Chadani et al., 2017). As the translation of MgtL aborts within the first six amino acid residues (MEPDPT) of MgtL, other proteins with MgtL-like N-terminal sequences could also induce premature translation abortion. We searched the 4,187 *E. coli* ORFs for those with sequences enriched in D, E and P in the six amino acid windows following the N-terminal methionine, and found ten genes as candidates. To determine whether these DE/P-enriched N-terminal sequences indeed cause IRD sequence-dependent translation discontinuation (hereafter referred to as “IRD”), we then fused the N-terminal ten amino acid sequences (Nt10) of the candidate genes to LacZ (Figure 1A) and measured the LacZ activities. As controls, we also constructed their “no DE/P” variants, in which the N-terminal D, E, and P residues were replaced by N (Asn), Q (Gln), and A (Ala), respectively.

**Figure 1.**
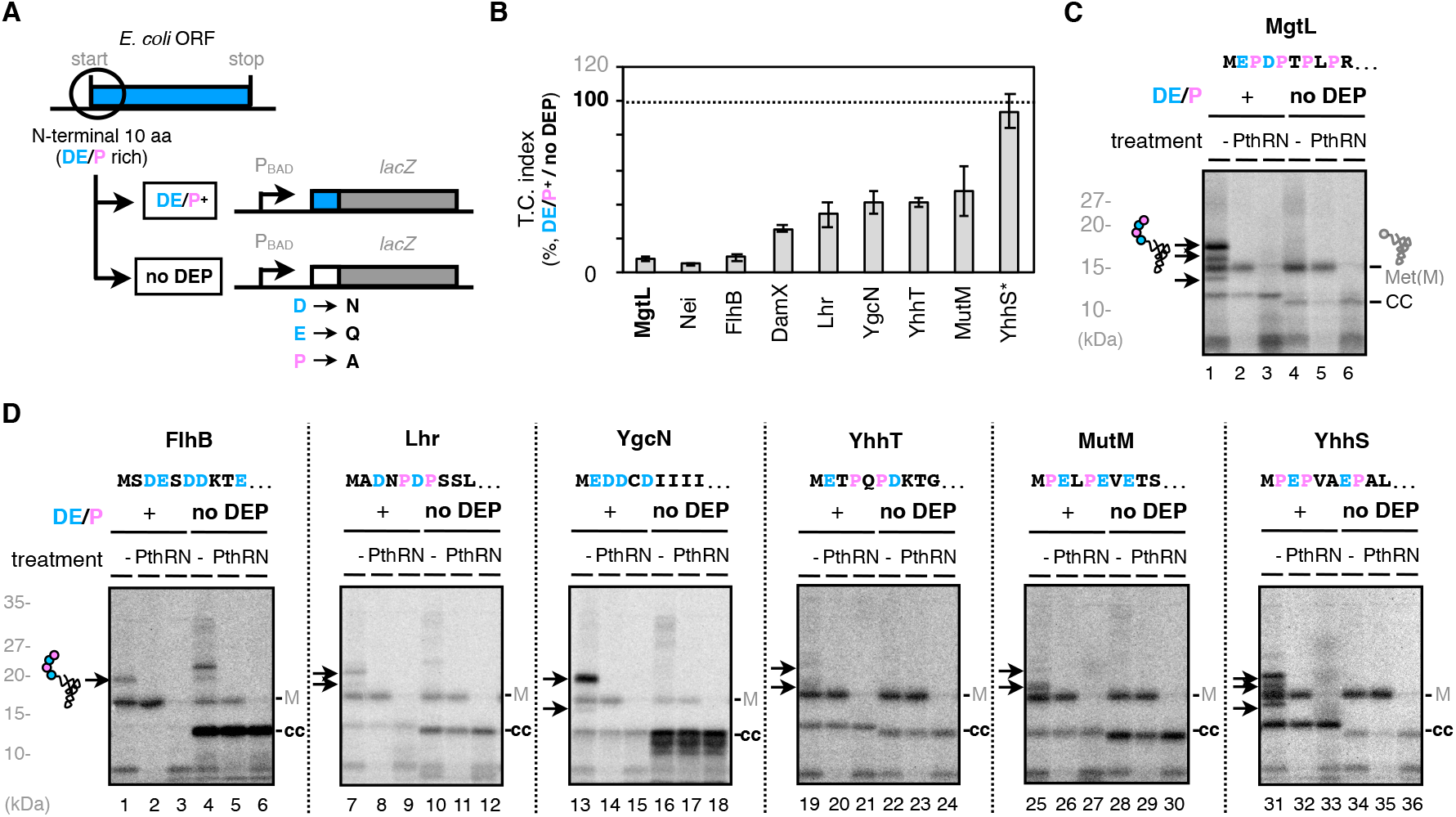
N-terminal DE/P clusters in *E. coli* ORFs induce IRD. A. The N-terminal 10 amino acids of the IRD-candidate ORFs are fused to *lacZ* (Nt10). To evaluate the translation continuity of the DE/P clusters in the ORFs (*DE/P^+^*), D, E and P in the N-terminal regions were replaced with N, Q and A, respectively (*no DEP*). B. N-terminal DE/P cluster-dependent translation attenuation *in vivo*. Each of the Nt10 constructs with or without DE/P residues were expressed in *E. coli*, and β-galactosidase activities were determined (Miller, 1972). The downstream translation frequency corresponding to IRD was calculated as the LacZ activity ratio [DE/P^+^ / no DEP], termed the translation continuation (T.C.) index. C. Peptidyl-tRNA accumulation during the *mgtL* translation. The Nt10 *mgtL*-*lacZ* construct and its variant with the DE/P replacement (no DEP) were translated by the PURE system in the presence of ^35^S-methionine, treated with peptidyl-tRNA hydrolase (Pth) as indicated, and separated by neutral pH SDS-PAGE with an optional RNase A (RN) pretreatment. Peptidyl-tRNAs are indicated by arrows with a schematic label. Radioactive formyl methionyl-tRNA and stop codon-terminated product are indicated as “M” and “CC” (translation-completed chain), respectively. D. Peptidyl-tRNA accumulation during the translation of other Nt10 constructs.

We normalized the enzyme activity of each construct by that of the respective no DE/P variant, to obtain the quantitative indications, termed the translation continuation (T.C.) index, in which 100% and 0% values represent no IRD and complete DE/P-dependent translation discontinuation, respectively. Out of the eight candidates with sufficient expression levels, seven showed translation continuation indices of less than 50 (Figure 1B). These translation anomalies at the N-terminal regions were independent of the 5’ untranslated regions (UTRs) (Figure S1A and S1B). In addition, a low Mg^2+^ concentration and the deletion of bL31, conditions that destabilize the 70S ribosome complex, enhanced the DE/P-dependent translation attenuation (Figure S1C and S1D). These results suggest that IRD occurred during the translation of the N-terminal DE/P-rich regions of these ORFs.

We next studied the translation of the Nt10 constructs in the PURE system, a reconstituted *in vitro* translation system (Shimizu et al., 2001), to recapitulate the translation abortion using defined translation components. Nt10 constructs with a shortened *lacZ* fragment were translated by the PURE system in the presence of ^35^S-methionine, and the synthesized products were separated by SDS-PAGE at neutral pH. The occurrence of IRD is indicated by the accumulation of peptidyl-tRNAs, which are sensitive to RNase A (RN, pre-electrophoresis treatment) or Pth (peptidyl-tRNA hydrolase, added to the translation mixture), but not puromycin (added to the reaction mixture). A typical result with MgtL is presented in Figure 1C, which shows the evident accumulation of a Pth-cleavable peptidyl-tRNA (*lanes* 1-3, arrows on *left*). As expected, the no DE/P variant of MgtL did not appreciably generate this product (*lanes* 4-6). The constructs bearing the other Nt10 sequences also accumulated Pth-cleavable peptidyl-tRNAs in DE/P-dependent manners (Figure 1D). Thus, the *mgtL*-like N-terminal DE/P-enriched sequences in the *E. coli* genes induce IRD in the reconstituted translation system.

### N-terminally adjacent nascent polypeptide segments in the exit tunnel counteract IRD in a length-dependent manner

The translation abortion by the N-terminal DE/P-enriched sequences (Figure 1) led us to investigate whether IRD occurs similarly when a DE/P sequence resides elsewhere, such as internal positions, in a polypeptide chain. To study IRD induction by internal DE/P motifs, we surveyed the *E. coli* genome and found 2,110 genes, out of the annotated 4,187 ORFs, that contain one or more DE/P-rich segment(s) (having four residues that are either D, E, or P in a six amino acid window). In most cases, the motifs were present at internal locations. We engineered the MgtL sequences with the EPDP sequence at various positions relative to the N-terminus, by inserting GFP-derived sequences between the initiator methionine and the EPDP motif. As shown in Figure 2A, the insertion of 238 amino acid residues counteracted the IRD-like translation anomalies at the repositioned EPDP motifs (Figure 2A).

**Figure 2.**
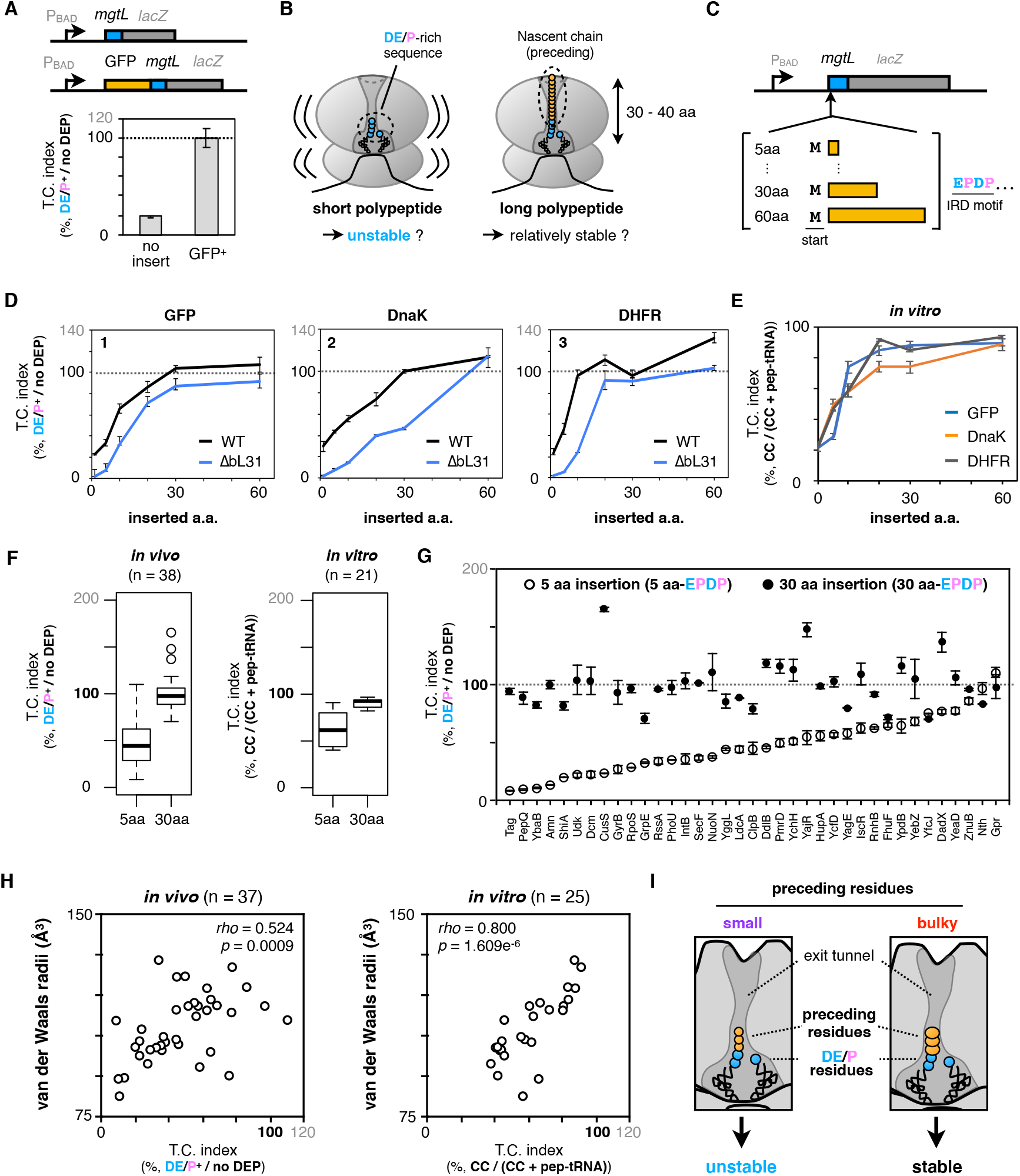
Exit tunnel-occupying nascent polypeptides minimize the risk of IRD in two different manners. A. IRD-dependent translation attenuation by *mgtL* at the early (*no insert*, upper schematic) or middle (*GFP^+^*, lower schematic) stage of translation elongation *in vivo.* Translation continuity indices were evaluated as in Figure 1. B. Schematic of the working hypothesis that a tunnel-occupying preceding nascent polypeptide stabilizes the translating ribosome to prevent IRD. C. Schematic of the *mgtL-lacZ* variants, in which various lengths of the N-terminal portions of GFP or other ORFs were inserted before the *mgtL* sequence. D. Inserted amino acid length-dependent IRD-counteraction. The counteracting effects were expressed as the downstream translation continuities [DE/P^+^/no DEP] of each *mgtL-lacZ* variant. The N-terminal regions of *E. coli* GFP (*panel 1*), DnaK (*panel 2*), or DHFR (*panel 3*) were utilized as the preceding nascent polypeptides. Results of wild type (*black line*) and IRD-prone ΔbL31 (*blue line*) cells are shown. E. IRD-counteracting effects of the N-terminal insertions in a reconstituted *in vitro* translation system (PURE system). The ratio of the completed chain (CC) per aborted peptidyl-tRNA (pep-tRNA) accumulated in the PURE system reaction was calculated as the downstream translation continuation of each *mgtL-lacZ* variant. F. IRD-counteracting effect of short and long nascent polypeptides within the tunnel. Downstream translation continuation of *mgtL-lacZ* variants with a 5 aa- or 30 aa-insertion (5 or 30 aa-EPDP) in the *in vivo* reporter (*left*, *n* = 37) or the *in vitro* translation assay (*right*, *n* =21) are represented by boxplots. G. Individual values of *in vivo* downstream translation continuation of the *mgtL-lacZ* variants with the 5 aa-(*white dots*) or 30 aa-EPDP constructs (*black dots*). H. Two-dimensional plots of downstream translation continuation of the 5 aa-EPDP constructs and averaged van der Waals molecular radii of inserted sequences. Both *in vivo* (*left*) and *in vitro* (*right*) experiments are represented with Spearman’s correlation coefficients. I. Schematic of the ribosomal tunnel occupancy by the bulkiness of the nascent polypeptide adjacent to the PTC.

When translating the wild-type *mgtL,* the ribosome accommodates at most a 6 amino acid-long nascent product in its exit tunnel before IRD occurs, leaving the tunnel almost unoccupied. By contrast, the exit tunnel of the ribosome translating the GFP-inserted variant of MgtL is fully occupied by the GFP part of the nascent chain when it is translating the repositioned EPDP motif. Thus, we hypothesized that the tunnel-spanning nascent polypeptide preceding the IRD sequence stabilizes the translating ribosome, and thus counteracts IRD (Figure 2B). We tested this hypothesis by inserting various lengths of sequences from a variety of origins before the EPDP motif of MgtL, as shown in Figure 2C. We chose various lengths of the N-terminal regions of GFP, *E. coli* DnaK and DHFR and evaluated their effects on the EPDP-mediated IRD in living cells. All of the nascent polypeptide segments preceding EPDP lowered the IRD efficiency in length-dependent manners (black line in Figure 2D). It is noteworthy that the IRD-counteracting effects of the inserted segments reached plateaus at a length of ~30 amino acids. Interestingly, this 30 amino acid length coincides with the length of the nascent polypeptide that spans the exit tunnel in an extended form (Jenni and Ban, 2003; Voss et al., 2006). When the same proteins were expressed in the bL31 deletion strain, the IRD effect was more pronounced than that in the wild-type host strain, and the protecting effects of the N-terminal polypeptide insertion became somewhat weaker. These results suggest that the possession of an upstream polypeptide sequence exerts an IRD-counteracting effect by stabilizing the translating ribosome. Consistent with this interpretation, we were able to reproduce the polypeptide insertion-dependent IRD alleviation by *in vitro* assays using the PURE system (Figure 2E), indicating that the IRD-counteraction does not require any factors other than the essential translation factors in this reconstituted system.

From these results, we propose that the nascent polypeptide chain within the ribosomal interior space generally contributes to the stability of the translating ribosome and the continuity of translation. To examine if any sequence specificity exists in the ribosome stabilizing ability of the tunnel-occupying polypeptide, we randomly chose 40 *E. coli* genes, and inserted 5 aa and 30 aa parts of their N-terminal regions into the *mgtL*-*lacZ* reporter, as described above. We successfully assessed the impacts of the sequences derived from 37 and 21 proteins by an *in vivo* reporter assay and *in vitro* translation, respectively (Figure 2F). As shown in the boxplots representation, the 30 aa insertions prevented IRD more effectively than the 5 aa insertions, both *in vivo* and *in vitro*. Thus, the polypeptide segments preceding the IRD sequence counteract the IRD in a largely sequence-independent manner, provided that they are long enough (~30 amino acids or longer) to span the exit tunnel.

### Bulky amino acids near the PTC also counteract IRD

We noticed that the 5 aa insertions had a rather broad spectrum of effects on IRD (Figure 2F). The individual *in vivo* data shown in Figure 2G indicate that the effects of the 5 aa insertions varied from nearly no effect to almost complete suppression of IRD (Figure 2G). A similar tendency was also observed in the PURE system analysis *in vitro* (Figure S2A). Thus, not only the nascent chain length but also some property of the amino acids in the immediate N-terminal vicinity of an IRD sequence can alleviate IRD, probably by stabilizing the 70S ribosome. We surveyed various properties of the inserted amino acids and found a positive correlation between the sum of the van der Waals radii of the five inserted amino acids and the IRD-counteracting effects *in vivo* and *in vitro* (Figure 2H). Residues with bulkier side chains prevented IRD more effectively than those with smaller ones (Figure 2I). By contrast, the average hydrophobicity showed a poor correlation with the effects (Figure S2B).

We tested the effects of bulky amino acids in the D-rich FLAG tag sequence (DYKDDDK), which exhibited significant IRD (Figure S2C). We mutated its YK part to pairs of identical amino acids, and found that the IRD propensities correlated with the bulkiness of the dipeptides at this position (Figure S2C and S2D). Taken together, bulky amino acid residues just preceding an IRD sequence counteract IRD. Our results suggest that the occupation of the exit tunnel by a nascent chain stabilizes the translating ribosome, by either the “length principle”, in which polypeptide chains with diverse sequences of ~30 residues can work, or the “size principle”, in which a short stretch of bulky residues can work.

### Multifaceted nascent chain-exit tunnel interactions contribute to the continuous translation elongation

Previous studies on ribosome arresting peptides revealed that nascent chains interact with the tunnel wall, particularly in the central constriction partially formed by the protein components uL4 and uL22 (Ito and Chiba, 2013) (Figure 3A). We addressed whether the mutations of these proteins affect the two modes of IRD antagonization studied above. We first focused on the role played by the uL22 loop (Figure 3B), and constructed mutant ribosomes with the mutant form lacking the β-hairpin loop (uL22Δloop; Figure 3B). We then developed a modified PURE system containing the mutant form of the ribosomes. The results revealed that the effects of the uL22Δloop mutation differ, depending on the preceding amino acid lengths. Whereas the uL22 mutated ribosome was almost indistinguishable from the wild-type ribosome in the IRD profile of the EPDP constructs with the upstream 5 amino acids (Figure 3C, Figure S3, *orange* line), the absence of the loop enhanced IRD when the upstream region contained 30 amino acids (Figure 3C, Figure S3).

**Figure 3.**
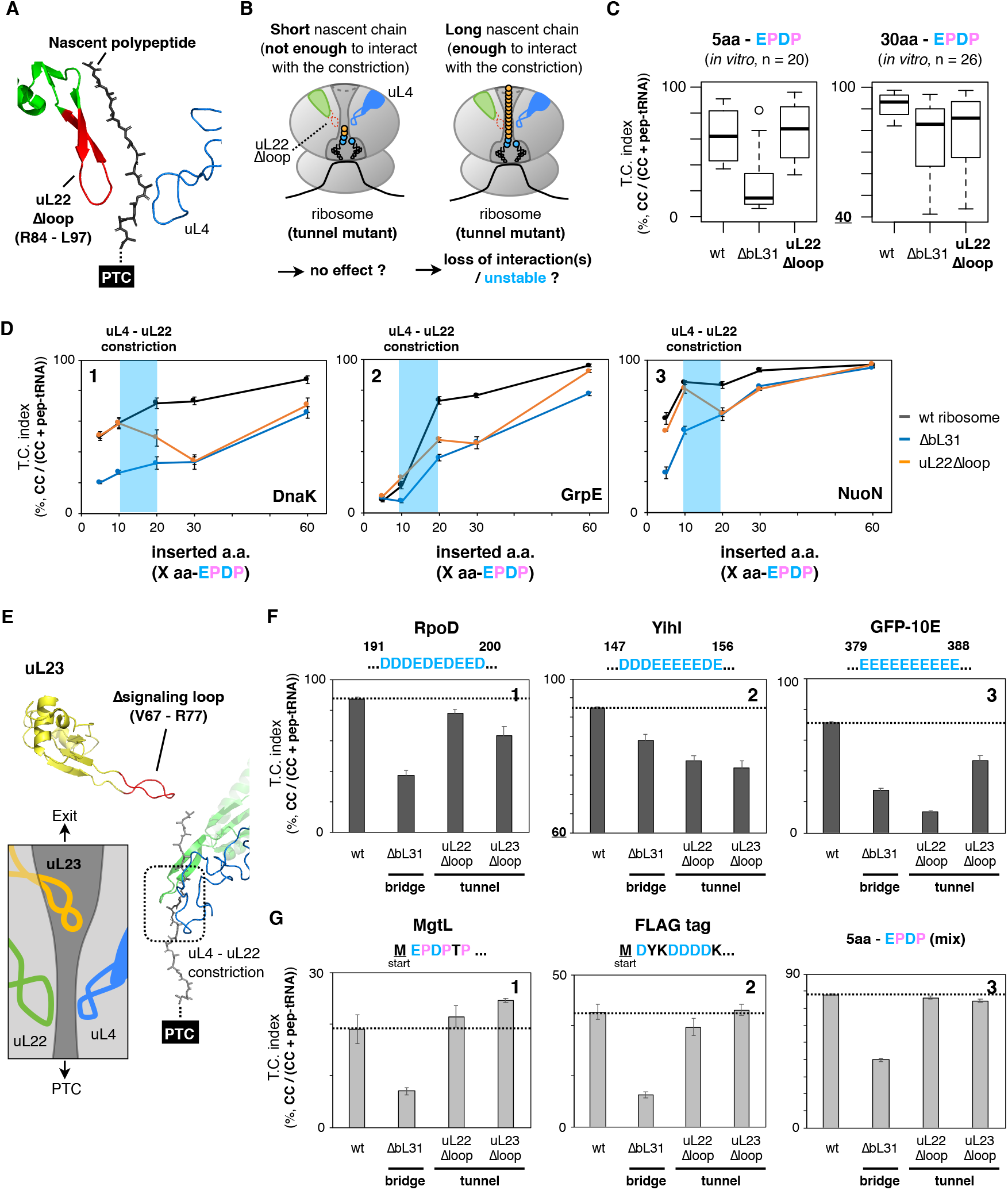
Effect of mutations in the constriction site within the exit tunnel on the IRD-counteraction. A. Structure of the uL4-uL22 constriction site (PDB: 4V5H). Deleted residues in the uL22Δloop mutation are indicated in red. B. Schematic of the working hypothesis that stabilization of the translating ribosome depends on interactions between the ribosomal tunnel and the tunnel-occupying nascent polypeptide. C. Inserted amino acid length-dependent IRD-counteracting effects in the uL22Δloop-ribosome. Messenger RNAs encoding a 5 aa-(*left*) or 30 aa-(*right*) insertion and EPDP (used in Figure 2) were individually translated by customized PURE systems including the indicated ribosome variants. Evaluated downstream translation continuation indices of wild type, ΔbL31 or uL22Δloop ribosomes are shown. D. Preceding nascent polypeptide-length dependency on the IRD counteraction in the ribosome variants. The N-terminal regions of *E. coli* DnaK (*panel 1*), GrpE (*panel 2*) or NuoN (*panel 3*) were utilized as preceding nascent polypeptides. E. Structure of the uL23 signaling loop (PDB: 4V5H) and schematic drawing of the tunnel structure. Deleted residues in the uL23Δloop mutation are indicated in red. F. Translation continuity of ORFs with the IRD-inducing motif located in the middle of the ORFs. Messenger RNAs encoding *rpoD* (*panel 1*), *yihI* (*panel 2*) or GFP-10E (*panel 3*) carrying 10 consecutive D/E residues were translated *in vitro* and the translation continuities were evaluated as described above. G. Translation continuity of ORFs with an IRD sequence without a preceding nascent polypeptide occupying the tunnel. Messenger RNAs encoding *mgtL* (*panel 1*), FLAG tag (*panel 2*) or a mixture of 5 aa-EPDP (*panel 3*) were translated *in vitro* and analyzed as described above.

This substrate-dependent effect of the uL22 mutation is in sharp contrast to our results with the ribosomes lacking bL31 (ΔbL31). The lack of bL31 invariably enhanced IRD (lowered the translation continuity), with both templates having the 5 aa and 30 aa insertions (Figure 3C. For individual data, see also Figure S3, compare *black* and *blue* lines). These results indicate that the β-hairpin loop of uL22 in the wild-type ribosome plays a role in counteracting IRD, but only in the presence of the tunnel-spanning upstream polypeptide region (Figure 3B). By contrast, bulky side chains near the PTC can stabilize the ribosome irrespective of the tunnel status (Figure 2I).

We further explored the relationships between the exit tunnel occupation and the stability of the translating ribosome, using various lengths of the N-terminal regions of DnaK, GrpE and NuoN that were attached N-terminally to the EPDP motif. The wild type and uL22Δloop ribosomes translated the EPDP constructs with the 5 amino acid- and the 10 amino acid-upstream regions with almost the same degrees of IRD, which differed depending on the sequences (Figure 3D, compare *black* and *orange* lines). However, the uL22Δloop ribosomes showed an increased IRD when 20 or more amino acids were present before the EPDP (Figure 3D, *orange* lines). Since previous studies suggested that the uL22 β-hairpin loop interacts with the nascent polypeptides at a 10-20 aa distance from the PTC (Bhushan et al., 2011; Seidelt et al., 2009; Shanmuganathan et al., 2019; Sohmen et al., 2015; Su et al., 2017), our data suggest that the interactions between the nascent polypeptide and the tunnel constriction site contribute to the elongation continuity beyond the IRD sequence, perhaps by changing the stability of the ribosome.

We next studied the effect of the other ribosomal tunnel mutation on IRD. We used uL23Δloop with the deletion of the uL23 signaling loop, which otherwise extends into the PTC-distal part of the exit tunnel (Bornemann et al., 2008, Figure 3E). We translated RpoD, YihI and the artificial GFP-10E (Chadani et al., 2017), which have acidic stretches starting from their 191^st^, 147^th^ and 379^th^ residues, respectively, using the PURE system with the mutated ribosomes (Figure 3F). The new substrate proteins with their acidic regions farther away from the N-termini exhibited rather weak IRD when translated with the wild-type ribosome, while the IRD was enhanced by the absence of bL31 (Figure 3F). As we observed with the uL22Δloop mutant, the deletion of the uL23 signaling loop enhanced IRD in all cases, suggesting that the uL23 loop in the wild-type ribosome contributes to the ribosome stability (Figure 3F). By contrast, the N-terminal IRD, without an insertion, was not affected by the tunnel mutations of the ribosome (Figure 3G). The latter results are expected, because these constructs are subject to translation discontinuation before accessing the uL22 and the uL23 tips in the tunnel (Figure 3B and 3E). These results suggest that nascent chains spanning the tunnel interact with the tunnel-exposed protein components of the ribosome to stabilize the translating ribosome and ensure translation continuity, irrespective of the amino acids being polymerized at the PTC. The involvement of multiple ribosomal proteins suggests that multiple interactions, which alone may be subtle, cumulatively contribute to the continuous translation.

### Proteome-wide amino acid distribution bias that minimizes the IRD risk

The features of the non-uniform translation revealed above might have affected the amino acid composition of proteins during the evolution of ORFs. Specifically, the N-terminal regions of ORFs might have avoided the enrichment of D, E, and P and preferred amino acid residues with bulky side chains to allow for robustly continuous translation elongation. We addressed this question by bioinformatics. The appearance of more than three D, E or P residues in ten-residue moving windows is lower for the N-terminal 20-30 positions of the *E. coli* ORFs than the following positions (Figure 4A). A previous bioinformatics analysis also showed a similar decline of negative charges at the beginning of the ORFs (Dao Duc and Song, 2018). We also observed that the van der Waals radii of amino acids in the N-terminal regions are larger than those in the other parts of ORFs (Figure 4B). Such biased amino acid composition features of the N-terminal regions are widely conserved among bacterial species, including *Bacillus subtilis, Staphylococcus aureus* and *Thermus thermophilus* (Figure S4). We suggest that the amino acid distribution biases were acquired during evolution as IRD-counteracting measures.

**Figure 4.**
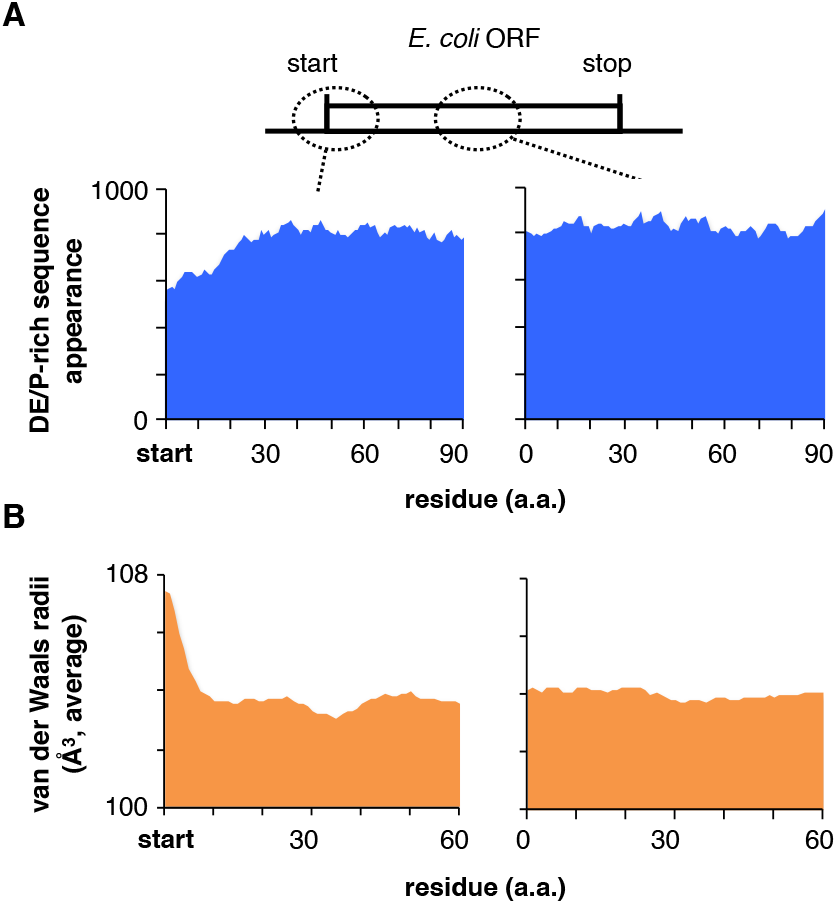
Genome-wide biases in amino acids distributions and van der Waals radii. A. Distributions of the DE/P-rich sequence (more than 3 DE/P in a 10 residue-moving window) appearance at the N-terminal regions (*left*) and in the middle (*right*) of entire *E. coli* ORFs. B. Averaged van der Waals molecular radii of every 5 aa-window at the N-terminal regions (*left*) and in the middle (*right*) of entire *E. coli* ORFs.

### Proteome-wide IRD and its counteraction revealed by ribosome profiling

In spite of the statistical tendency revealed above, the N-terminal regions of ORFs still have many DE/P- or small amino acid-rich regions (Figure 4), with some of the former inducing IRD (Figure 1). To capture IRD globally, we performed ribosome profiling (Ribo-seq, Ingolia et al., 2009) and monitored the ribosomal occupancy along mRNAs. We expected that the translation of ORFs with a DE/P-enriched sequence in the N-terminal regions would cause a decline in the ribosome occupancy (the number of ribosome-protected RNA fragments, RPFs) in the following 3’ region, by the stochastic release of the ribosomes from the mRNA due to IRD (Figure 5A). To survey ORFs with the above-mentioned characteristic patterns of ribosome distribution, we compared the sum of the RPF counts at the first 10 codons with those at the following 10 codons (Figure 5B). We first focused on *ybeX* as one of the ORFs that exhibited the above mentioned Ribo-seq patterns. The N-terminal segment (MSDDNSHSSDT) of *ybeX* is enriched in Asp and small-sized amino acids (Ser and Thr). The RPFs from the *ybeX* mRNA decreased after the 11^th^ codon (Figure 5C). A biochemical analysis showed that a peptidyl-tRNA accumulated in an Asp-dependent manner during the *in vitro* translation of *ybeX* (Figure S5A), substantiating the occurrence of an elongation cessation.

**Figure 5.**
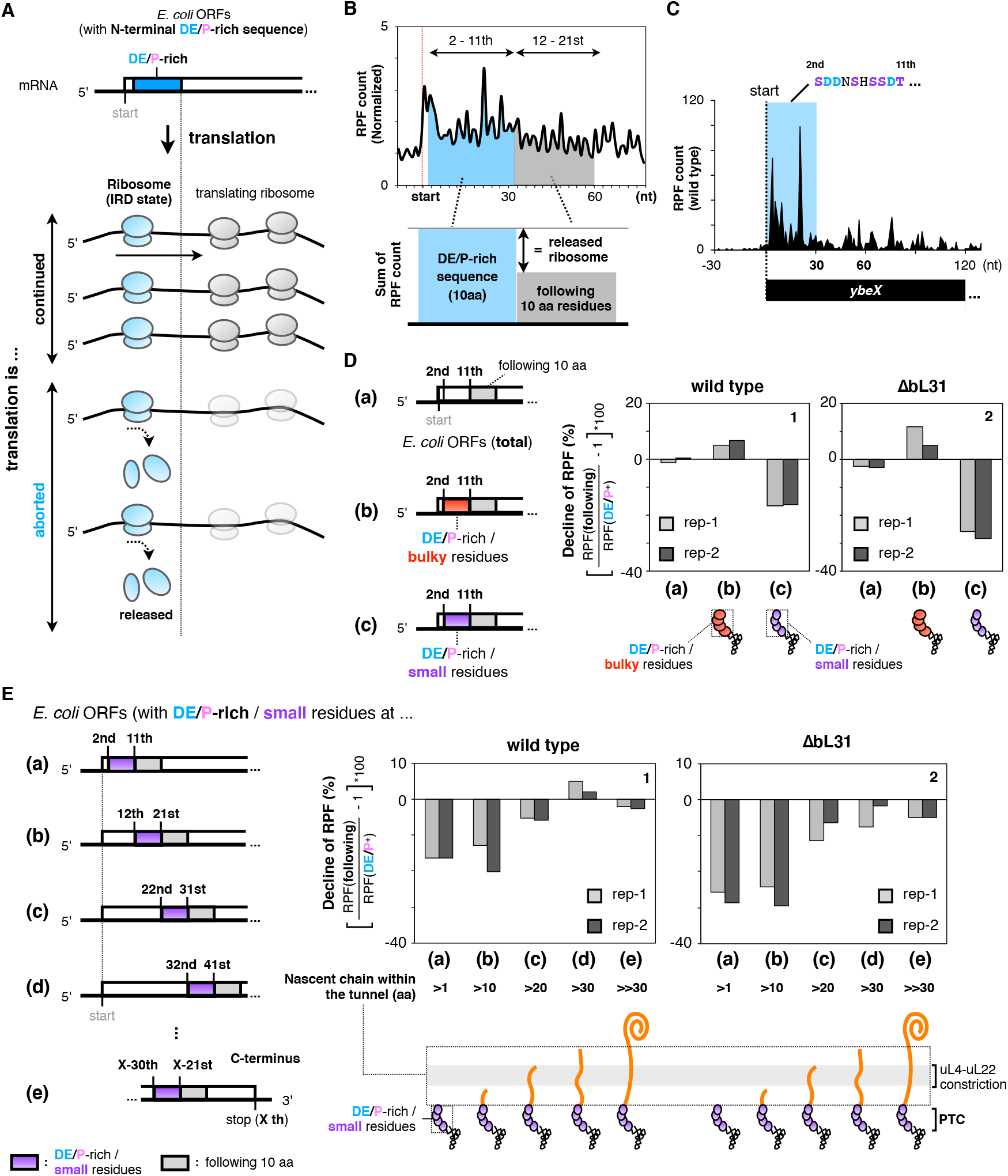
Ribosome profiling analysis to survey the ribosome occupancy around the N-terminal regions. A. Schematic representation showing that the ribosome is released from the mRNA by IRD, leading to a decline in ribosome-protected fragments (RPFs) after translation of IRD-inducing sequences. B. IRD frequency around the N-terminal region was evaluated from the ratio of RPF counts in the 12^th^ −21^st^ aa to those in the 2^nd^ −11^th^ aa including DE/P-rich residues. C. Distribution of the RPFs on the *ybeX* mRNA in wild type cells. The region of the N-terminal acidic-rich sequence is highlighted. D. Ratio of RPF counts in the 12^th^-21^st^ aa to those in the 2^nd^-11^th^ aa. The data are classified as follows: Total: all dataset (a), *E. coli* ORFs with DE/P-rich and bulky (b) or small (c) radii-residues in N-terminal regions. A detailed representation of the classification is shown in Supplementary Figure S5B. The data from two biological replicates of wild type (*panel 1*) or IRD-prone ΔbL31 (*panel 2*) cells are shown. Schematics at the bottom represent the nascent chain occupying the tunnel during the translation of the 2^nd^-11^th^ aa. E. Ratio of RPF counts in 10 amino acids with “small” and DE/P-rich residues to those in the following 10 amino acids at different stages of elongation. RPF counts for 10 amino acids with “small” and DE/P-rich residues were summed from the most N-terminal (2^nd^ −11^th^, (a)) to the subsequent every 10 amino acids windows, (b)-(d). RPF counts for 10 amino acids with “small” and DE/P-rich residues in (e) were summed from X-30^th^ to X-21^st^, where X represents the last amino acid in the ORFs. The data from two biological replicates of wild type (*panel 1*) or IRD-prone ΔbL31 (*panel 2*) cells are shown. The corresponding data for “bulky” and DE/P-rich residues are shown in Supplementary Figure S5C.

We next analyzed the global Ribo-seq data to compare the sum of the RPF counts at the first 10 codons with those at the following 10 codons. We extracted the ORFs enriched in DE/P residues in their N-terminal regions, and then classified them further into those enriched in small amino acids and those enriched in bulky amino acids (Figure S5B). The analysis revealed that ~16% of RPFs were lost after 10 codons (30 nucleotides) for ORFs with an abundance of small-sized amino acids in their N-terminal regions (Figure 5D, panel 1). The bL31-deleted cells showed more pronounced decreases in RPFs after the 10^th^ codon (Figure 5D, panel 2), further supporting our contention that the global low-occupancy of the ribosomes after the risky sequences represents IRD. By contrast, we did not observe any decrease in the RPF for ORFs containing DE/P-rich and large-sized amino acids in the N-terminal region (Figure 5D), consistent with our findings that bulky amino acids antagonize IRD.

We next undertook a more systematic survey of the N-terminal regions of *E. coli* proteins for the occurrence of IRD-like phenomena. In this analysis, we searched for sequences enriched in DE/P and small-sized residues in the successive 10 amino acid windows moving from the N- to C-terminal regions, as schematically shown in Figure 5E (a)-(e). We then analyzed the numbers of RPFs across the putative IRD sequences. We found 15-20% declines when the windows starting at the second or the 12^th^ codons were examined (Figure 5E, panel 1 (a), (b)). Such tendencies were gradually lost for windows moving further toward the 3’ direction, in which the elongating nascent polypeptide should have possessed successively longer sequences preceding the IRD determinant. The results with the windows of 22-31 (c), 32-41 (d) and at the C-terminus (e) indicated that the ribosomes essentially continue to translate these regions even when they encounter DE/P- and smaller residue-enriched segments. The bL31 deletion uniformly enhanced the decline of RPFs, again supporting that the elongation discontinuations were due to IRD (Figure 5E, panel 2). We did not observe equivalent decreases in RPFs when we extracted the DE/P-enriched sequences that were associated with bulky amino acid residues (Figure S5C), suggesting that the nascent chain “length”- and amino acids “bulkiness”-dependent mechanisms synergistically affect the translation continuity. Collectively, our Ribo-seq analysis revealed that DE/P-enriched segments that also contained small-sized residues generally induce IRD when they are present at N-terminal regions.

### Tunnel-occupying nascent polypeptides stabilize the translating 70S ribosome by bridging the 50S and the 30S subunits

Our findings that the nascent polypeptide spanning the exit tunnel counteracts the IRD-mediated translation discontinuation strongly suggested that the tunnel-occupying nascent polypeptide functions as an inter-subunit bridge, like the bL31-supported B1b peripheral bridge. We performed a sucrose gradient centrifugation analysis to assess the stability of the 70S ribosome, with or without the nascent polypeptide within the tunnel. As reported previously (Chadani et al., 2017; Fischer et al., 2015; Lilleorg et al., 2017; Ueta et al., 2017), the deletion of bL31 destabilizes the 70S complex structure, resulting in partial dissociation into the 30S and 50S subunits in cell lysates (Figure 6A). We found that the small amount of the 70S complex remaining in the ΔbL31 cell lysate was completely eliminated by a treatment with puromycin. This drug cleaves peptidyl-tRNA to produce peptidyl-puromycin, which neither interacts with mRNA nor remains ribosome-associated (*blue line*). By contrast, chloramphenicol, which blocks translation elongation, preserved the 70S complex (*red line*). These results are consistent with the notion that the nascent polypeptide stabilizes the 70S structure.

**Figure 6.**
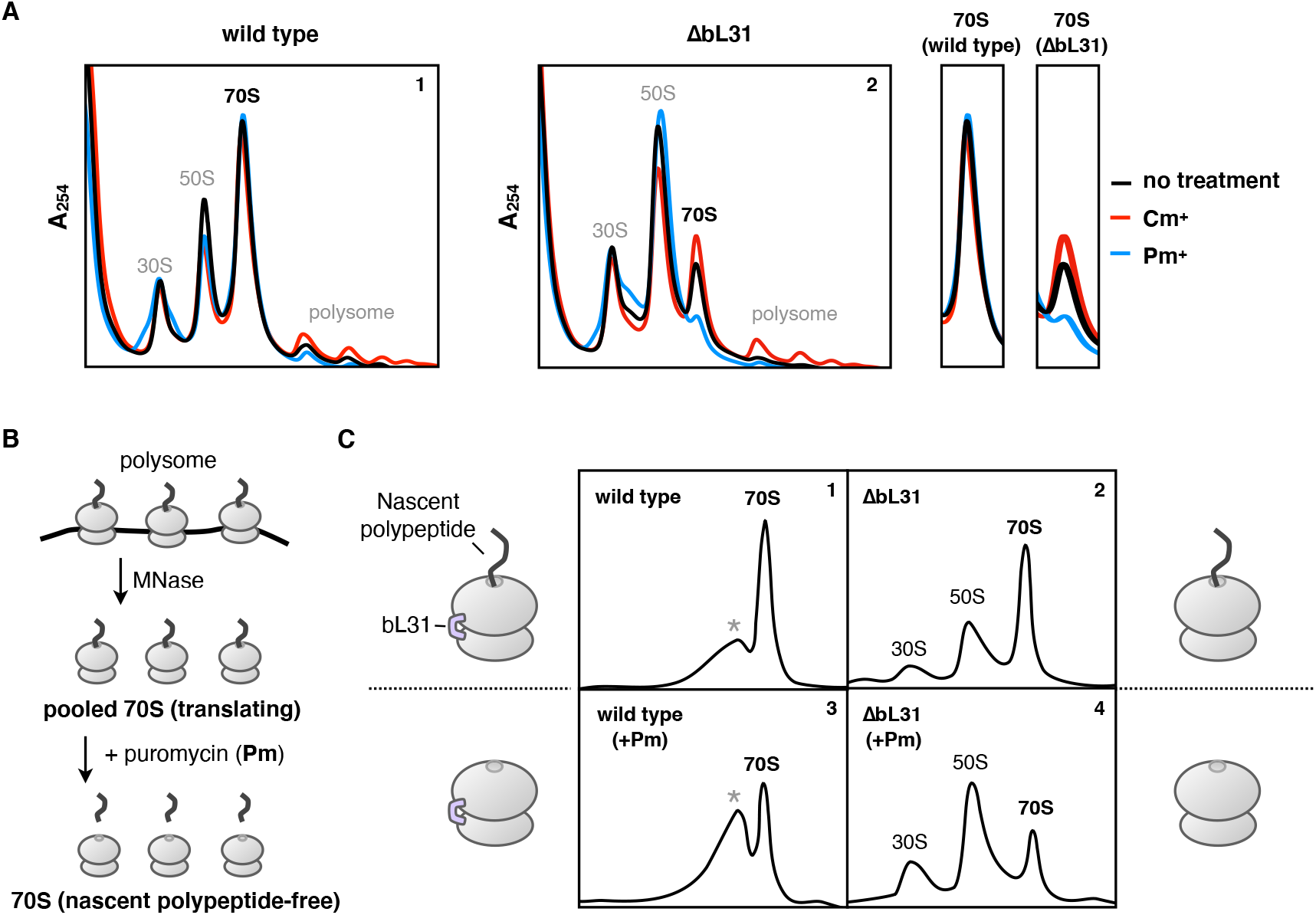
Exit tunnel-occupying nascent polypeptide physically stabilizes translating 70S ribosome as an inter-subunit bridge. A. Stability of the translating ribosome complex with or without the bL31 protein. Cellular extracts of wild type (panel 1) or ΔbL31 (panel 2) cells were fractionated by sucrose density gradient ultracentrifugation (SDG) in the presence of 10 mM magnesium ion (no treatment, *black line*). Prior to the extraction, *E. coli* cells were treated with puromycin (Pm, *blue line*) or chloramphenicol (Cm, *red line*) to remove or keep the nascent polypeptide within the ribosome, respectively. Distributions of the ribosomes were monitored by A_254_ measurements. Fractions including the 70S ribosome are expanded for comparison. B. Experimental design to evaluate the stability of the translating ribosomes. MNase: micrococcal nuclease. C. Stability of the 70S ribosome complex with or without a tunnel-occupying nascent polypeptide in wild type (panels 1 and 3) or ΔbL31 (panels 2 and 4) cells. Pooled 70S translating ribosomes with (panels 3 and 4) or without (panels 1 and 2) the puromycin treatment were fractionated by SDG in the presence of 10 mM magnesium ion. Recorded A_254_ values are shown. The peaks indicated as the asterisk (*) in panel 1 and 3 seem to be an aberrant form of the 70S ribosome due to the MNase treatment.

To substantiate the concept that the nascent polypeptide stabilizes the association of the large and small ribosomal subunits more directly, we compared the stabilities of the 70S ribosomes with a tunnel-traversing nascent polypeptidyl-tRNA and those that had lost the intact polypeptidyl-tRNA. We isolated the 70S ribosomes in the process of translation by treating a polysome preparation with micrococcal nuclease, which cleaves unoccupied mRNA. The resulting nascent chain-ribosome complexes exhibited the ~70S sedimentation speed. We then treated the isolated 70S complex with puromycin and centrifuged the samples again (Figure 6B). Wild type ribosomes sedimented at ~70S irrespective of the puromycin treatment (Figure 6C, panels 1 and 3). By contrast, the bL31-lacking ribosomes were dissociated by the puromycin treatment into the 30S and 50S subunits (Figure 6C, panel 4). Thus, the nascent polypeptides (peptidyl-tRNAs) within the exit tunnel stabilize the translating 70S ribosome complex even in the absence of the bL31 bridge. It should be noted that the nascent polypeptides within the 70S ribosomes in this experiment consisted of a mixture of endogenous gene products, which do not necessarily contain IRD sequences around the PTC.

## Discussion

Recent advances in translation studies have revealed the widespread occurrence of noncanonical translation progression (Dever et al., 2020; Samatova et al., 2021). We showed that elongation pausing is frequent in *E. coli,* by capturing intermediate molecules with tRNAs at their ends (Chadani et al., 2016). More recently, we discovered the IRD phenomena (Chadani et al., 2017), in which the translation of certain acidic sequences, sometimes containing a proline, leads to the destabilization of the 70S ribosome and translation cessation without involving a stop codon. In the present study, we addressed the generality, the position effects, and the sequence context-dependence of potentially IRD-inducing DE/P-rich motifs in *E. coli*, featuring both model constructs and the native genes in the genome. Our results showed that nascent polypeptides in the interior of the translating ribosome have regulatory roles over the stability of the engaged ribosomes. In particular, we found that the nascent chain exerts stabilizing effects on the 70S structure, thus compensating for the potential IRD induction by a risky DE/P-rich part in the following region of the same polypeptide.

In the translocation step of translation, the large and small subunits undergo a relative rotation, which their association must withstand to produce a protein product (Bock et al., 2015; Horan and Noller, 2007; Voorhees and Ramakrishnan, 2013). Thus, the ribosome is equipped with multiple bridges, including bL31, that tether the large and small subunits in the functional monosome (Chadani et al., 2017; Fischer et al., 2015; Lilleorg et al., 2017; Ueta et al., 2017). We have shown that the growing nascent chain itself also contributes to the stabilization of the protein synthesizing machine. Although we initially found the stabilization effect of nascent chains from their counteraction to IRD, our sedimentation analysis using cell lysates (Figure 6) suggested that endogenous sources of nascent polypeptidyl-tRNA, not necessarily combined with a DE/P sequence, stabilize the 70S structure. Thus, we propose that the ribosome stabilization by the tunnel-occupying nascent chain represents a general life principle.

Earlier studies reported the peptidyl-tRNA-dependent stabilization of the 70S structure under Mg^2+^-depleted conditions (Duin et al., 1970; Kohler et al., 1968; Ron et al., 1968; Schlessinger et al., 1967). Other reports described the destabilization of a nascent chain-free ribosome complex under high ionic conditions (Blobel and Sabatini, 1971; Zylber and Penman, 1970). Whereas these observations complement our findings, they were made before the discovery of the ribosomal exit tunnel. In addition, cryptic and apparently risky sequences like the IRD-motifs were unknown, and thus the physiological significance of the early findings remained unexplored. Our work can therefore be regarded as a revisitation of this problem with the cutting-edge knowledge of translation dynamics. Our results have engendered the emergence of the novel concept that the nascent products of translation, in conjunction with the exit tunnel, contribute to the stability of the ribosome in action. We named it *<u>T</u>*unnel-occupying nascent *<u>P</u>*olypeptide-dependent *<u>S</u>*tabilization (TPS). TPS enables the ribosome to be a universal translation machine that can synthesize polypeptides of any sequence without discontinuation to produce the life-supporting proteome of the species. The C-terminal end of a nascent polypeptide is covalently tethered to a tRNA that interacts with the 30S component and the mRNA in the 30S decoding center. Meanwhile, its more N-terminal region resides in the 50S subunit, entering into and occupying the exit tunnel. Thus, a nascent polypeptide bridges the small and large subunits. TPS represents a self-stabilization mechanism to prevent the dissociation of the translating ribosome.

Despite the potentially deleterious outcomes, dozens of *E. coli* genes with N-terminal DE/P-rich sequences still evoke IRD. Why do these genes keep IRD sequences, which destabilize the ribosome? One possibility is that cells harness IRD as a regulatory scheme at the translational and/or transcriptional levels. A typical example might be found in the translation of MgtL, which serves as a Mg^2+^ sensor for regulation of the MgtA Mg^2+^ transporter (Chadani et al., 2017). Our present ribosome profiling and biochemical experiments revealed that the expression of *ybeX* (*corC,* Giménez-Mascarell et al., 2019), a putative Mg^2+^/Co^2+^ efflux pump, can be repressed in an IRD-dependent manner. An increased Mg^2+^ concentration in the cell would stabilize the ribosome and decrease the IRD-mediated premature translation cessation, leading to the enhancement of the *ybeX* expression and the consequent Mg^2+^ efflux. It is also conceivable that the N-terminal IRD sequences are advantageous for cells to respond to certain stresses other than changes in the Mg^2+^ concentration that affect the ribosome stability (Rasouly and Ron, 2009). The N-terminal IRD sequences might be a hidden code for the “ribosome stability regulon” that reshapes the proteome upon emergency conditions.

Nascent polypeptides preceding the DE/P-rich sequences antagonize IRD. In the “length”-dependent but sequence-independent mode of the stabilization, the nascent chain might act as an intersubunit scaffold. The deletion of either the uL22 β-hairpin loop or the uL23 signaling loop weakens the effect but does not completely abolish it, suggesting that multiple weak interactions between the interior of the tunnel and the nascent polypeptide contribute to the stabilization. The diverse effects of the ribosomal mutations affecting different ribosomal proteins, such as uL22 and uL23, imply that the ribosome and the nascent chain communicate with each other by the local diversities, producing cumulative weak interactions to perform the biological role of the “peptidyl-tRNA bridge”, although such interactions poorly inhibit the nascent chain movement through the tunnel.

These non-uniform features of the interactions remind us of the interactions between the tunnel and ribosome arresting peptides (RAPs, Ito and Chiba, 2013). Each part of the tunnel architecture differently interacts with each of the RAP sequences to arrest translation. For example, the alternation of either of uL22 Gly91, Ala93 or the 23S rRNA A2058 residue attenuates the translational arrest induced by RAP in *E. coli* SecM, but not that of *E. coli* TnaC (Cruz-Vera et al., 2005; Nakatogawa and Ito, 2002). In this respect, RAPs are similar to the peptides involved in TPS, but differ in that RAPs require specific polypeptide sequences at a particular site within the tunnel. However, too many specific interactions between the tunnel and the nascent polypeptide would inhibit its passage through the tunnel, resulting in translational arrest events. From this viewpoint, TPS and translation arrest share the same principle, but with different strengths. TPS promises the continuity of translation, but too much stabilization might be one of the driving forces of translational arrest. The widespread occurrence of translational pausing during the synthesis of the *E. coli* proteome might reflect such interactions within the tunnel (Chadani et al., 2016). Since the translational arrest events have physiological impacts, modulation of TPS could influence the elongation velocity, leading to cotranslational control of gene expression.

The bulkiness, defined as the van der Waals radii, of the amino acids in the nascent chain adjacent to PTC also counteracted IRD. It is likely that the “bulkiness”-dependent stabilization plays a more important role during the early stage of translation, at which the “length” works poorly. The “length”-dependent mechanism apparently lacks a preference for amino acid sequences, but its effect would be insufficient until the nascent polypeptide fulfills the tunnel (no less than 20 aa). In contrast, the “bulkiness”-dependent stabilization necessitating larger residues restricts the usage of amino acids, unlike the “length”-dependent one. We assume that these two stabilization mechanisms complementarily compensate for their respective insufficiencies to avoid nonproductive premature termination. The strong biases on disfavored DE/P-rich sequences and favored bulky amino acids in the N-terminal regions of bacterial ORFs are in agreement with this assumption (Figure 4 and Figure S4). Many previous studies have pointed out that the sequence features just adjacent to the start codon significantly influence the efficiency of protein biosynthesis (Allert et al., 2010; Goodman et al., 2013; Kudla et al., 2009; Osterman et al., 2020; Salis et al., 2009; Sprengart et al., 1996; Verma et al., 2019). Considering the “bulkiness” of the nascent polypeptide when interpreting the datasets might disclose new landscapes from previous studies.

The “size (bulkiness)”-dependence indicates the relationship between the geometry of the tunnel structure and the stabilization mechanism. Lu and colleagues reported the “size”-dependent rearrangement of the nascent polypeptide at the eukaryotic uL4-uL22 constriction site, indicating that larger residues have more interaction chances than smaller ones (Lu et al., 2011). The translation of consecutive tryptophan residues, the largest natural amino acid, inhibits elongation by the eukaryotic ribosome, in agreement with this observation (Dimitrova et al., 2009). The prokaryotic uL4-uL22 constriction site is slightly broader than that of eukaryotes (Dao Duc et al., 2019), and tryptophan repeats do not induce translational anomalies in *E. coli* (Chadani et al., 2017). However, the prokaryotic tunnel forms another constriction site at the entrance of the tunnel, which is as narrow as the eukaryotic uL4-uL22 constriction site (tunnel radii are below 4 Å, Dao Duc et al., 2019; Fischer et al., 2015). Taken together with these facts, we assume that the squeezing of bulky residues by this “entry constriction site” represses the unwinding of the growing products around the PTC, thus stabilizing the 70S complex. Other bacterial ribosomes show similar architectural characteristics, supporting this assumption.

Finally, this study provides an answer to the naive question of why the ribosome has an exit tunnel for the nascent polypeptide. A tunnel may not be needed, if the essential function of the ribosome is simply to polymerize the polypeptides. Stabilization of the translating ribosome by preparing the tunnel for occupation with its own product would be a positive feedback system to continue the elongation. There might be a trade-off between the stabilization effects and the obstacles of translation elongation. The tunnel architecture, which is less than 30~40 aa in length, might represent a compromise for these characteristics.

The schemes of the two stabilizing mechanisms provide a glimpse of the molecular details of IRD. It is likely that “length”- and “size (bulkiness)”-dependent stabilization commonly minimizes the fluctuations of polypeptidyl-tRNAs within the ribosome complex. Taken together with the putative negative-negative repulsion between the D/E residues and the rRNA component of the tunnel wall (Nissen et al., 2000), we assume that IRD interrupts the correct positioning of the polypeptidyl-tRNA, resulting in a disruption of the codon-anticodon interactions and the following subunit dissociation. Understanding such aspects of the ribosomes will clarify the basis of ribosomal functions and the secondary genetic codes hidden within amino acid sequences.

## Author Contributions

YC and NS performed experiments; YC, NS, TN, YI, SI and HT conceived the study, designed experiments and analyzed the results; YC and HT supervised the entire project; YC and HT wrote the manuscript.

## Acknowledgments

We thank Koreaki Ito for critical reading and editing of the manuscript, Eri Uemura for technical support, the Bio-support Center at Tokyo Tech for DNA sequencing, and The National BioResource Project, *E. coli* (National Institute of Genetics, Japan) for providing the ASKA library clones and Keio collection strains. This work was supported by MEXT Grants-in-Aid for Scientific Research (Grant Numbers JP26116002, JP18H03984, JP20H05925 to HT, JP17H05679, JP17H04998, JP19K22406, JP20H05784 to SI, JP17K15062, JP19K16038 to YC), and a grant from the Ohsumi Frontier Science Foundation to YC.

## STAR Methods

### CONTACT FOR REAGENT AND RESOURCE SHARING

Please direct any requests for further information or reagents to the lead contact, Hideki Taguchi.

## EXPERIMENTAL MODEL AND SUBJECT DETAILS

### *E. coli* strains, plasmids, and primers

*E. coli* strains, plasmids and oligonucleotides used in this study are listed in Table S1. The ASKA plasmid library (Kitagawa et al., 2006) were obtained from National Institute of Genetics (NIG). Plasmids were constructed using standard cloning procedures and Gibson assembly. Detailed schemes were summarized in Table S1, and sequences of constructed plasmids will be uploaded to the Mendeley repository.

PCR-amplified DNA fragment from pDGT1 using primers PR0058 and PR0059 was electroporated into strain BW25113 harboring pKD46 (Datsenko and Wanner, 2000) and pCY927 (*rpsJ-rpsQ* operon), selecting for a trimethoprim (20 μg/ml)-resistant transformant. Genome-introduced Δ*rpsJ*-*rpsQ*::FRT-DHFR-FRT mutation was transferred by Phage P1 into MG1655 harboring pCY873 (*rpsJ-rpsQ* operon), pCY920 (*rpsJ-rpsQ* operon with uL22Δloop mutation), or pCY1457 (*rpsJ-rpsQ* operon with uL23Δloop mutation), respectively.

## METHODS DETAILS

### In vitro translation and product analysis

A coupled transcription-translation reaction was performed using PUREfrex 1.0 (GeneFrontier) in the presence of ^35^S-methionine at 37°C for 30 min. DNA templates were prepared by PCR, as summarized in Table S2. Mutant ribosomes (ΔbL31, uL22Δloop, uL23Δloop) purified as described (Ohashi et al., 2007) were used when indicated. After 30 min incubation, the reaction mixture was treated with 200 μg/ml of puromycin for 5 min at 37°C. Half of the mixture was further treated with 1 μM of purified Pth for 10 min at 37°C when indicated. The reaction was stopped by dilution into an excessive volume of 5% TCA. After standing on ice for 10 min or more, samples were centrifuged for 3 min at 4°C, and the supernatant was discarded by aspiration. Precipitates were then vortexed with 0.9 ml of acetone, centrifuged again, and dissolved in SDS sample buffer (62.5 mM Tris-HCl pH 6.8, 2% SDS, 10% glycerol, 50 mM DTT) that had been treated with RNasecure (Ambion). Finally, the sample without Pth treatment was divided into two portions, one of which was incubated with 50 μg /ml of RNase A (Roche) at 37°C for 30 min and separated by WIDE range SDS-PAGE system (Nakalai tesque).

### Translation continuation (TC) index *in vitro*

The ratio of the translation-completed chain (CC) against Pth-sensitive polypeptidyl-tRNAs (pep-tRNA), which signifies the occurrence of IRD-induced abortion in the total *in vitro* translation products, was calculated as translation continuation (TC) index. In detail, the radioactivity (^35^S-methionine) proportion of “CC” and “pep-tRNA” among the sample with neither Pth- nor RNase-treatment were quantified by Multi Gauge (Fujifilm), and the TC value (in %) is obtained by the following formula.

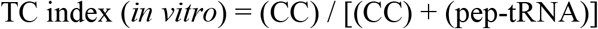

The average of three independent experiments was presented as TC indices, except for Figure 3C (Figure S3).

### β-galactosidase assay

#### TC index *in vivo*

*E. coli* cells harboring *lacZ* reporter plasmid were grown overnight at 37°C in LB medium supplemented 100 μg/ml ampicillin. They were inoculated into fresh LB medium containing 2× 10^−4^ % arabinose and 100 μg/ml ampicillin and were grown at 37°C for ~2.5 hrs (A_660_ = ~0.4). Then 20 μl portion was subjected to β-galactosidase assay as described (Miller, 1972). Estimated miller unit (*m.u.*) were subjected to the following formula to calculate the TC index (*in vivo*).

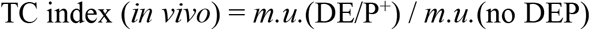

TC indices were calculated from three independent experiments, and plasmids used in each experiment were summarized in Table S2.

#### Mg^2+^-depletion

*E. coli* cells harboring *lacZ* reporter plasmid were grown overnight at 37°C in M9 medium (Miller, 1972) supplemented with 0.1% casamino acids, 0.1 mM CaCl_2_, 50 μM MgSO_4_, 50 μg/ml of thiamine, 0.4% glycerol and 100 μg/ml ampicillin. They were inoculated into fresh M9 medium containing 2× 10^−4^ % arabinose, and were grown at 37°C for 4 hrs. Then, 20 μl portion was subjected to β-galactosidase assay as described (Miller, 1972).

The concentration of MgSO_4_ in M9 medium was adjusted to 10 mM or 10 μM when indicated.

### Amino acids distribution analysis

We calculated the number of ORFs with more than three D,E and P residues within 10-amino acids moving windows in both N-terminal and internal regions in entire *E. coli* ORFs. Datasets of protein sequences were obtained from UniProt (https://www.uniprot.org).

### Ribosome profiling

#### a. Sequencing of ribosome-protected fragment (RPF)

*E. coli* MG1655 (wild type) or ECY0228 (MG1655 ΔbL31) strain was grown overnight in LB medium at 37°C. They were inoculated into fresh LB medium and were grown at 37°C. When A_660_ reached to 0.5, cells were treated with 100 μg/ml of chloramphenicol and collected by filtration (0.45 μm MF membrane). Filter-trapped *E. coli* cells were scraped and dropped into liquid nitrogen dispensed to 50 ml corning tube in advance. In addition, RP buffer (20 mM Tris-HCl pH 7.5, 150 mM NH_4_Cl, 10 mM MgCl_2_, 1 mM DTT, 5 mM CaCl_2_, 1% Triton X-100, 100 μg/ml chloramphenicol) was added to form frozen droplets. The cell droplets were disrupted by Multi-Beads Shocker [MB601U(S), Yasui Kikai], thawed and treated with 25 unit/μl of Turbo™ DNase (Thermo Fisher Scientific) for 10 min on ice. After centrifugation at 20,000 g for 10 min at 4 °C, the supernatant was collected and 35 μg RNA per sample was treated with 150 unit of Micrococcal Nuclease (MNase, Roche) for 45 min at 25°C. RNA fragments ranging from 17 to 50 nt were gel-excised as ribosome-protected fragment (RPF). Following library preparation was performed as previously described (McGlincy and Ingolia, 2017). A Ribo-Zero rRNA Removal Kit for Bacteria (Illumina) was used to deplete rRNA contamination. The libraries were sequenced on a HiSeq4000 (Illumina). After depleting the reads originating from noncoding RNAs, the remaining reads were mapped to the *E. coli* genome sequence (NC_000913.2). Empirically, we defined an A-site position, which is essentially the same as the 3′ assignment as described previously (Woolstenhulme et al., 2015).

#### b. Classification of *E. coli* ORFs by sequential features

“DE/P-rich” sequence

We defined two or more D, E or P residues within continuous three codons as “DE/P^+^ cluster”. Furthermore, two or more “DE/P^+^ clusters” within the continuous ten codons were defined as “DE/P-rich” sequences that potentially induce IRD and utilized for classification of *E. coli* ORFs.

“Van der Waals radii”

The averaged van der Waals radii of ten residues belonging to “DE/P-rich” sequence at indicated region was calculated and utilized for classification of *E. coli* ORFs as follows.

“DE/P-rich sequence” with bulky-sized residues: 40% of the larger one

“DE/P-rich sequence” with small-sized residues: 40% of the smaller one

#### c. Calculation of normalized RPF

We utilized house-made script to describe the distribution of normalized RPF counts among the subsets of *E. coli* ORFs classified as described above. Briefly, the relative RPF count in each nucleotide position was calculated for each ORF at first. Then, the obtained relative RPF count in each position was averaged for each subset. We excluded the ORFs with less than 150 bp length, poor expression (less than 400 RPF count/gene) or concentrated RPF count at specific site (b0621, b2094, b2218, b3709 and b2744) to convince the statistical analyses. The sum of normalized RPF counts among the indicated region was compared to that of following 10 amino acids residues to estimate the frequency of IRD-dependent translation abortion.

### Polysome profile

*E. coli* cells were grown in LB medium overnight at 37°C. They were inoculated into fresh LB medium and were grown at 37°C. When A_660_ reaches to 0.5, cells were optionally treated with 20 μg/ml puromycin for 10 min or 200 μg/ml chloramphenicol for 1 min. Cell culture was mixed with crushed ice and centrifuged at 8,000 rpm for 3 min at 4°C. Pelleted cells were resuspended by polysome buffer [20 mM Tris-acetate pH 7.5, 150 mM NH_4_Cl, 10 mM Mg(OAc)_2_, 1 mM DTT, (200 μg/ml chloramphenicol was added when indicated)] and dropped into liquid nitrogen dispensed to 50 ml corning tube in advance. Frozen cell droplets were disrupted by Multi-Beads Shocker [MB601U(S), Yasui Kikai] and treated with 10 unit of Turbo™ DNase (Thermo Fisher Scientific) for 20 min on ice. Lysate was centrifuged at 20,000 g for 10 min at 4°C. The supernatant was layered onto the top of 10-30% sucrose gradient containing the polysome buffer in Open Top Polyclear™ Centrifuge Tubes (14 × 89 mm, SETON) and centrifuged at 39,000 rpm for 4 h at 4°C (Beckman OptimaL-90K, SW41-Ti), followed by fractionation using Gradient Station (BIOCOMP) equipped with MICRO COLLECTOR AC-5700 (ATTO). Distribution of ribosome was monitored by A_254_ measurements using BIO MINI UV MONITOR (AC-5200S, ATTO).

### Stability assay for 70S translating ribosome complex

*E. coli* cells were grown in LB medium overnight at 37°C. They were inoculated into 2x YTPG medium supplemented with Antifoam SI (Fujifilm-WAKO) till A_660_ reaches to 3.0. Cells were collected and extracted as described in polysome profile analysis without antibiotic treatment. Prepared cell extract was fractionated by 10-50 % sucrose gradient ultracentrifugation at 39,000 rpm for 2.5 h at 4°C (Beckman OptimaL-90K, SW41-Ti). The fractions containing polysome (larger than the 70S monosome) were collected and again centrifuged at 50,000 rpm for 3 h at 4°C (Beckmann Optima™ TLX Ultracentrifuge, TLA100.3) to pellet down the polysome particles. The pellet was once rinsed by the polysome buffer and resolved by overnight-standing within the polysome buffer supplemented with 5 mM CaCl_2_ at 4 °C. Fifty μl of prepared polysome (A_260_ = ~8.0) was treated with 100 μg/ml of puromycin for 1 h at room temperature (24°C) with gentle mixing in the presence of 300 unit of Micrococcal Nuclease (MNase, Roche). Reaction mixture was mixed with 100 μl of polysome buffer and layered onto the top of 10-30% sucrose gradient containing the polysome buffer in Open Top Polyclear™ Centrifuge Tubes (14 × 89 mm, SETON) and centrifuged at 39,000 rpm for 3 h at 4°C (Beckman OptimaL-90K, SW41-Ti) to record A_254_ as described above.

### Data analyses

Statistical analyses were conducted by using the software R (https://www.r-project.org).

## Notes

### Competing Interest Statement

The authors have declared no competing interest.

