## Supplementary information for "Nascent polypeptide within the exit tunnel ensures continuous translation elongation by stabilizing the translating ribosome"

**Supplementary Figures S1-S5**

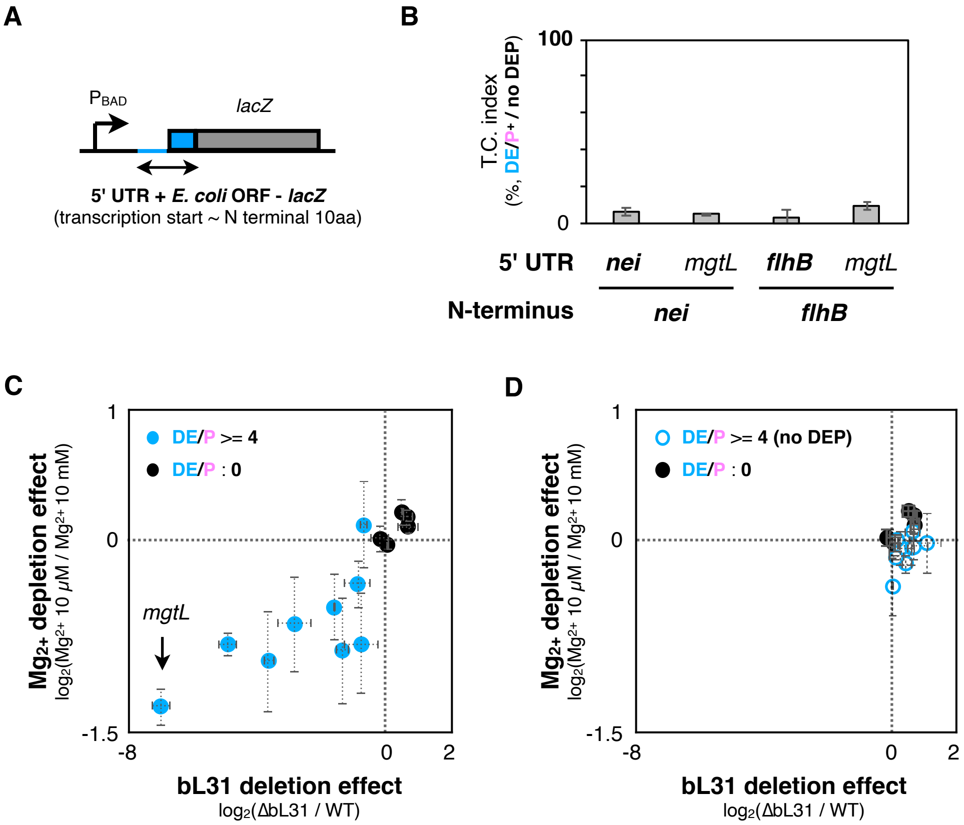


**Supplementary Figure S1**

**A** and **B**. 5' untranslated region (UTR) contributes little to the N-terminal IRD events. Schematic of the DNA template to examine the contribution of 5' UTR for translation attenuation by the N-terminal DE/P-rich residues (**A**), and downstream translation continuation (T.C.) evaluated by *in vivo* reporter assay (**B**).

**C** and **D**. Two-dimensional plots of the effects on $Mg^{2+}$-depletion (low $Mg^{2+}$/high $Mg^{2+}$) and the bL31 deletion (bL31/ WT) calculated from the LacZ activity ratios. In ***C***, the effects for the DE/P-rich Nt10 candidates (filled *blue* circles) and those for ORFs carrying the DE/P-free N-terminal regions (filled *black* circles) are shown. In ***D***, the DE/P residues in the Nt10 ORFs were replaced by NQ/A, and then the DE/P-free Nt10 ORFs (open *blue* circles) were examined for the translation continuation assay.

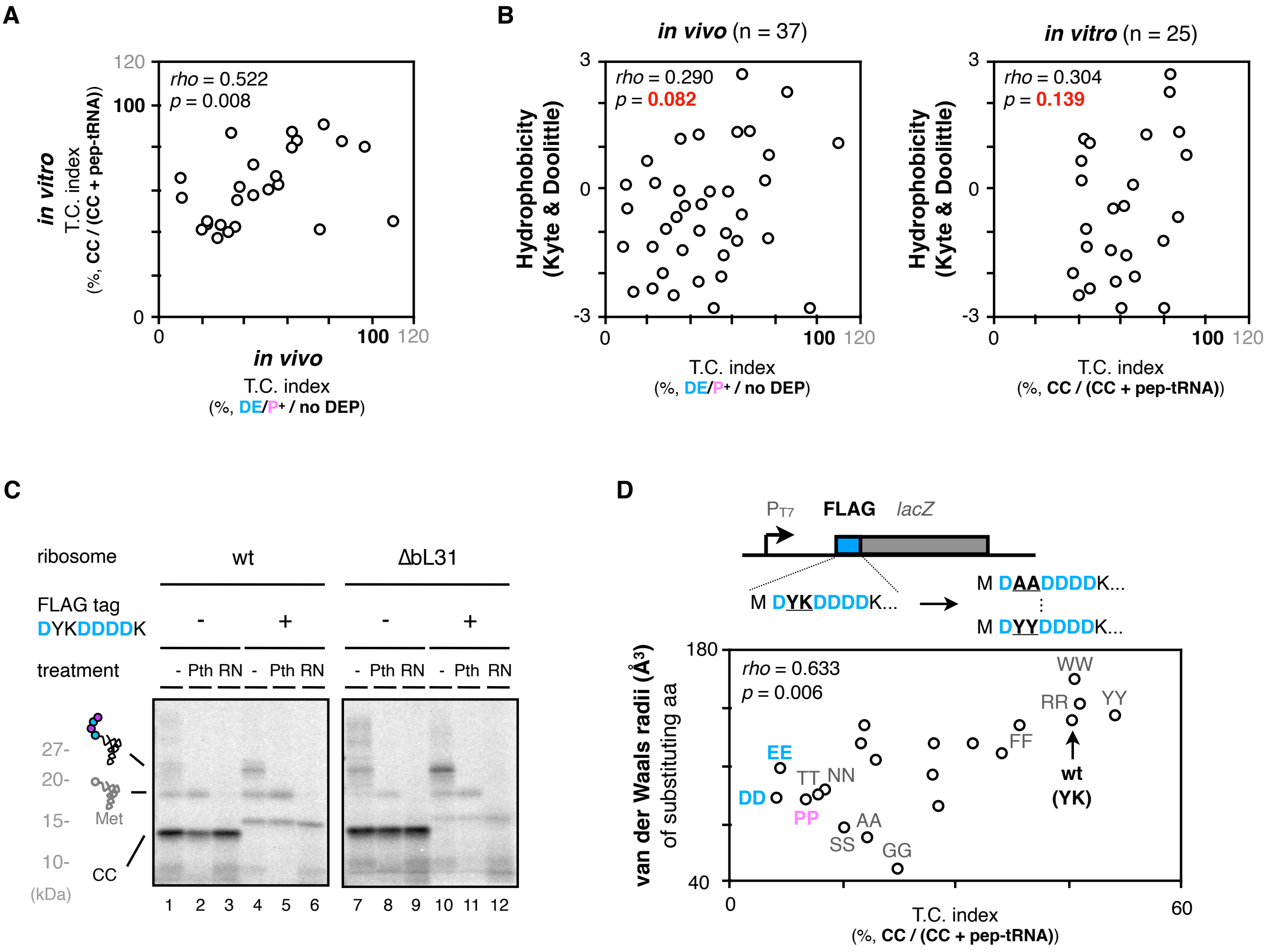


**Supplementary Figure S2**

**A.** Correlation between *in vivo* reporter and *in vitro* translation assays. *In vivo* and *in vitro* results of the 5 aa-EPDP constructs in Figure 2H were compared by two-dimensional plot. Spearman's correlation coefficients are also shown.

**B.** Two-dimensional plots of downstream translation continuation of the 5 aa-EPDP constructs and the average hydrophobicity of inserted amino acid sequences. Results of *in vivo* (*left*) and *in vitro* results (*right*) were shown.

**C.** Translation of the N-terminal FLAG tag-*lacZ* construct induces the accumulation of abortive peptidyl-tRNAs. *In vitro* translation was performed using standard (lane 1-6) or a customized PURE system equipped with ΔbL31 ribosome (lane 7-12) in the presence of $^{35}$S-methionine, and polypeptide products were separated by neutral pH SDS-PAGE.

**D.** Dependency of the molecular radii of amino acids on the translation continuation using FLAG tag as a model IRD-prone sequence. Second tyrosine and third lysine residues of the N-terminal FLAG tag-*lacZ* were replaced with 20 kinds of dipeptide motif as described in the upper schematic. FLAG tag variants were translated *in vitro*, and the correlation between the downstream translation continuation and the van der Waals molecular radii of substituted amino acids are plotted in two-dimensional. The result of Spearman's correlation test is also shown.

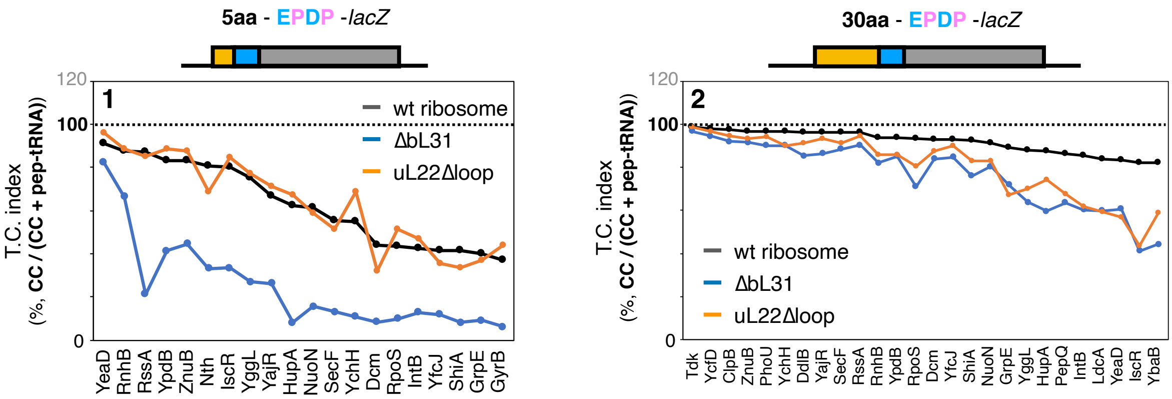


**Supplementary Figure S3**

Inserted amino acids length-dependent IRD-counteracting effects in the uL22Δloop-ribosome. Messenger RNA coding 5 aa- (*panel 1*) or 30 aa (*panel 2*) insertion and the EPDP used in Figure 2G were individually translated by customized PURE systems, including ribosome variants as indicated. Individual translation continuation indices of wild type (*black line*), ΔbL31 (*blue line*), or uL22Δloop ribosomes (*orange line*) are shown.

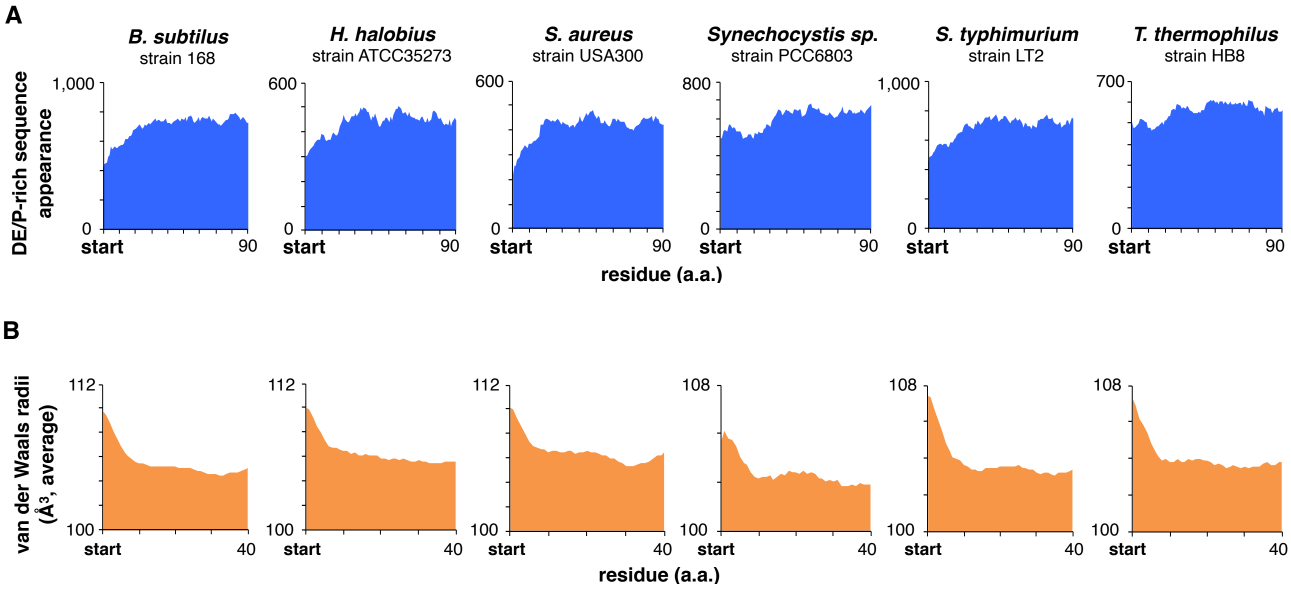


**Supplementary Figure S4**

**A** and **B**. Appearance of the DE/P-rich sequences (**A**) or average van der Waals molecular radii of 5 aa-moving window (**B**) at the N-terminal regions in the total ORFs of various bacterial species.

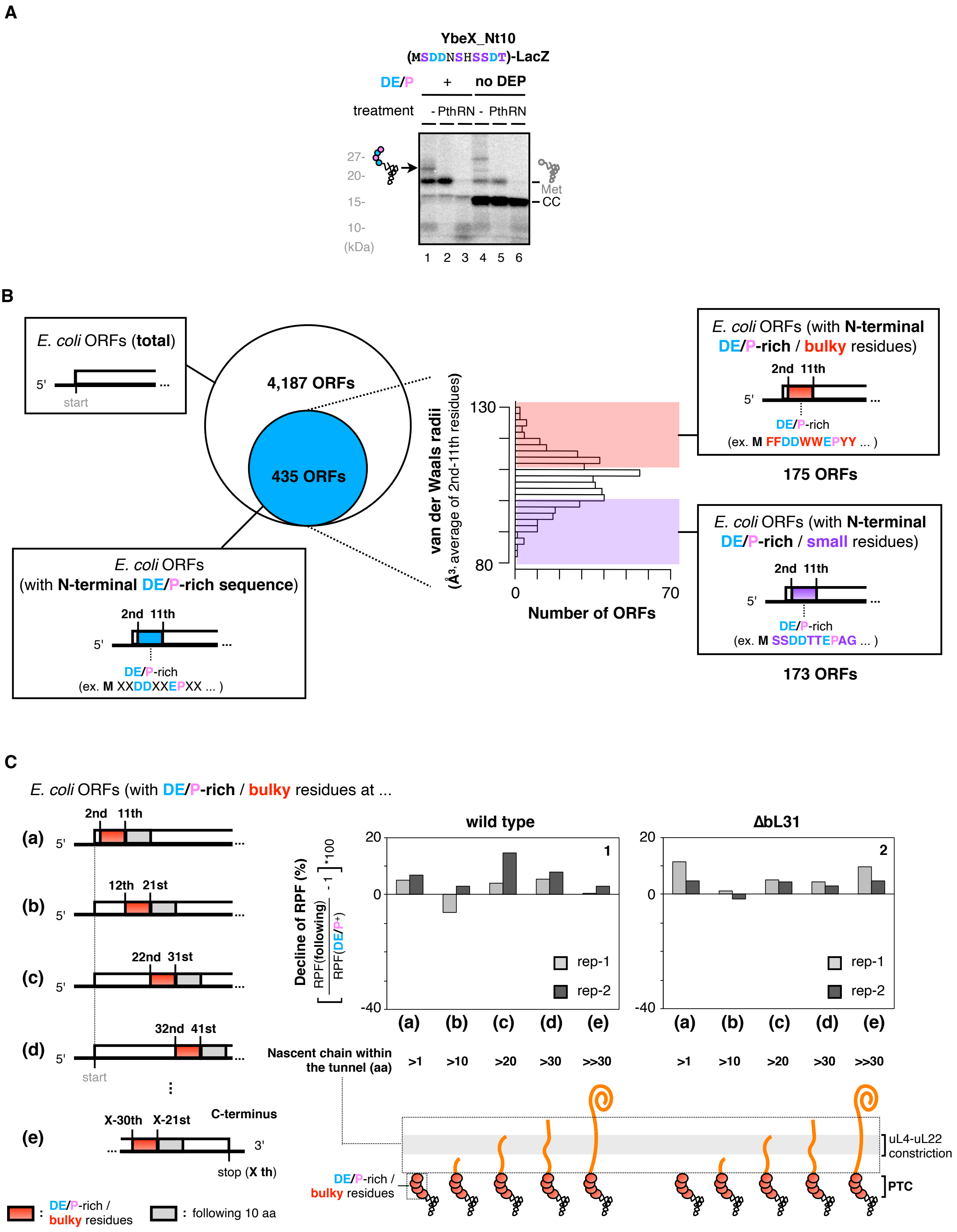
A
YbeX_Nt10
(MSDDNSHSSDT)-LacZ
DE/P
+
no DEP
treatment
- PthRN - PthRN
27-
20-
15-
10-
(kDa)
Met
CC
1 2 3 4 5 6
B
E. coli ORFs (total)
5'
start
4,187 ORFs
435 ORFs
E. coli ORFs
(with N-terminal DE/P-rich sequence)
2nd 11th
DE/P-rich
(ex. M XXDDXXEPXX ... )
van der Waals radii
(Å3, average of 2nd-11th residues)
130
80
0
70
Number of ORFs
E. coli ORFs (with N-terminal DE/P-rich / bulky residues)
2nd 11th
DE/P-rich
(ex. M FFDDWWEPYY ... )
175 ORFs
E. coli ORFs (with N-terminal DE/P-rich / small residues)
2nd 11th
DE/P-rich
(ex. M SSDDTTEPAG ... )
173 ORFs
C
E. coli ORFs (with DE/P-rich / bulky residues at ...
(a) 2nd 11th
(b) 12th 21st
(c) 22nd 31st
(d) 32nd 41st
start
(e) X-30th X-21st C-terminus
3'
stop (X th)
: DE/P-rich / bulky residues
: following 10 aa
wild type
ΔbL31
Decline of RPF (%)
[RPF(following) / RPF(DE/P+) - 1]*100
20
0
-40
1
2
rep-1
rep-2
(a) (b) (c) (d) (e)
Nascent chain within the tunnel (aa)
>1 >10 >20 >30 >>30
uL4-uL22 constriction
PTC
DE/P-rich / bulky residues

**Supplementary Figure S5**

**A.** Peptidyl-tRNA accumulation during the *ybeX* translation. *In vitro* translation products of *ybeX*_Nt10-*lacZ* or its "no DEP" variant were separated by SDS-PAGE with optional treatments as indicated.

**B.** Classification of *E. coli* ORFs by the N-terminal amino acid (aa) sequences in terms of the DE/P-richness and the bulkiness. At first, ORFs carrying DE/P-rich sequences ($n$ = 435) at 2-11$^{th}$ aa were extracted from the entire ORFs [$n$ = 4,187, (a) in Figure 5D]. They were further classified as DE/P-rich/bulky [$n$ = 175, (b) in Figure 5D] and DE/P-rich/small [$n$ = 173, (c) in Figure 5D] ORFs according to the average van der Waals radii of 2-11$^{th}$ aa.

**C.** Ratio of RPF counts in 10 amino acids with "bulky" and the DE/P-rich residues to those in following 10 amino acids at the different stages of elongation. The RPF counts for 10 amino acids with the "bulky" and the DE/P-rich residues were summed from the most N-terminal (2$^{nd}$ -11$^{th}$, (a)) to subsequent every 10 amino acids windows, (b)-(d). The RPF counts for 10 amino acids with the "bulky" and the DE/P-rich residues in (e) were summed from X-30$^{th}$ to X-21$^{st}$, where X represents the last amino acids in the ORFs. The data from two biological replicates of wild type (*panel 1*) or the IRD-prone ΔbL31 cells (*panel 2*) are shown.
